# International travelers and genomics uncover a ‘hidden’ Zika outbreak

**DOI:** 10.1101/496901

**Authors:** Nathan D. Grubaugh, Sharada Saraf, Karthik Gangavarapu, Alexander Watts, Amanda L. Tan, Rachel J. Oidtman, Jason T. Ladner, Glenn Oliveira, Nathaniel L. Matteson, Moritz U.G. Kraemer, Chantal B.F. Vogels, Aaron Hentoff, Deepit Bhatia, Danielle Stanek, Blake Scott, Vanessa Landis, Ian Stryker, Marshall R. Cone, Edgar W. Kopp, Andrew C. Cannons, Lea Heberlein-Larson, Stephen White, Leah D. Gillis, Michael J. Ricciardi, Jaclyn Kwal, Paola K. Lichtenberger, Diogo M. Magnani, David I. Watkins, Gustavo Palacios, Davidson H. Hamer, for the GeoSentinel Surveillance Network, Lauren M. Gardner, T. Alex Perkins, Guy Baele, Kamran Khan, Andrea Morrison, Sharon Isern, Scott F. Michael, Kristian G. Andersen

**Author notes:** These authors contributed equally to this work (Nathan D. Grubaugh, Sharada Saraf, Karthik Gangavarapu). These authors jointly supervised this work (Andrea Morrison, Sharon Isern, Scott F. Michael, Kristian G. Andersen). Correspondence and requests for materials should be addressed to N.D.G. or S.F.M.

## Abstract

The ongoing Zika epidemic in the Americas has challenged public health surveillance, response, and control systems. Even as the epidemic appears to be near its end in the Americas, it is unclear whether substantial Zika virus transmission may still be ongoing. This issue is exacerbated by large discrepancies in local case reporting and significant delays in detecting outbreaks due to surveillance gaps. To uncover locations with lingering outbreaks in the Americas, we investigated travel-associated Zika cases diagnosed in the United States and Europe to identify signatures of transmission dynamics that were not captured by local reporting. We found that a large and unreported Zika outbreak occurred in Cuba during 2017, a year after peak transmission in neighboring countries, with cases still appearing in 2018. By sequencing Zika virus from infected travelers, we show that the 2017 outbreak in Cuba was sparked by long-lived lineages of Zika virus introduced from multiple places in the Americas a year prior. Our data suggest that while aggressive mosquito control in Cuba may initially have been effective at mitigating Zika virus transmission, in the absence of vaccines, herd immunity, or strong international coordination, such control measures may need to be maintained to be effective. Our study highlights how Zika virus may still be ‘silently’ spreading in the Americas and provides a framework for more accurately understanding outbreak dynamics.

## Introduction

The recent Zika epidemic in the Americas is a testament to how rapidly mosquito-borne viruses can emerge and spread, and has revealed flaws in our surveillance and response systems (Grubaugh et al., 2018a; Morens and Fauci, 2017). Due, in part, to high rates of subclinical infections and overlapping symptoms with infections from dengue and chikungunya viruses (Mitchell et al., 2018), Zika virus was circulating for more than a year and a half before it was first detected in Brazil (Faria et al., 2017). By the time Zika virus was discovered in May of 2015 (Zanluca et al., 2015) and recognized for its ability to cause severe congenital disease (França et al., 2016; Mlakar et al., 2016), the virus had already spread from Brazil to more than 40 countries (Faria et al., 2017; Grubaugh et al., 2017; Metsky et al., 2017; Thézé et al., 2018). By mid 2017, reports from the Pan American Health Organization (PAHO, 2017a) revealed Zika virus activity throughout the Americas was waning, prompting predictions for the end of the epidemic (e.g. (O’Reilly et al., 2018)) and the removal of the World Health Organization’s (WHO) “Public Health Emergency of International Concern” status (WHO, 2016a, 2016b).

Coordinated response efforts during the early stages of the Zika epidemic were ultimately contingent on countries detecting cases and reporting them to international health agencies (Lessler et al., 2016), including PAHO. For Zika virus and other *Aedes aegypti* mosquito-borne viruses - including dengue and chikungunya viruses - that disproportionately impact those with limited resources (Braga et al., 2010; Gardner et al., 2018; Netto et al., 2017), accurate local reporting is especially problematic. Not only are people in poor living conditions more likely to be exposed to infected mosquitoes, but such communities often have less access to adequate healthcare, resulting in more cases going undetected (Hotez, 2016; LaBeaud, 2008). Pockets of virus transmission that occur in countries with inadequate reporting can therefore facilitate ‘hidden’ outbreaks, increasing the risk of infected travelers causing outbreaks in new regions of the world. Thus, underreported or unrecognized local outbreaks may prolong epidemics, and hinder global efforts aimed at halting virus spread.

Infectious disease surveillance of international travelers has long been an effective method to detect pathogens circulating in resource-limited areas (Hamer et al., 2017; Harvey et al., 2013; Leder et al., 2013, 2017; Wilder-Smith et al., 2012). We hypothesized that similar frameworks could be leveraged to improve Zika virus surveillance, as many regions in the Americas affected by the epidemic attract large volumes of international visitors from countries with stronger healthcare and surveillance systems (Wilder-Smith et al., 2018). In this study, we used international travel data, coupled with virus genomics, to detect ongoing Zika virus transmission that was missed by local reporting. We discovered a large Zika outbreak in Cuba that was not reported to PAHO (PAHO, 2017a) or other public health agencies, and thus went undetected to the international community. We show that the outbreak in Cuba peaked in 2017, when the epidemic in the rest of the Americas was waning (PAHO, 2017a), and estimate that it was as large as those in neighboring countries that occured the year before. By sequencing Zika virus from infected travelers, we also show that the outbreak in Cuba was caused by multiple introductions from elsewhere in the Caribbean and Central America. Overall, our study outlines a framework for how traveler surveillance data, combined with virus genomics, can detect ‘hidden’ outbreaks and reconstruct transmission dynamics when local data are insufficient.

## Results

### Uncovering an unreported Zika outbreak in Cuba

Tracking the spread of epidemics requires accurate case reporting and strong international collaboration. Failure to do so leaves us vulnerable to surveillance ‘blind spots’, with the potential for prolonging epidemics and increasing their geographic spread. Zika virus was first detected in Brazil in May, 2015 (Zanluca et al., 2015), yet studies have shown that the epidemic started at least one and a half years prior to its discovery (Faria et al., 2017). Early in 2016, 48 countries across the Americas reported local outbreaks (PAHO, 2017b), with case numbers peaking later that year. By mid 2017, new Zika cases were no longer being reported to the international community (PAHO, 2017b). Due to widespread surveillance gaps and inconsistent reporting, however, we hypothesized that local Zika outbreaks could still be occurring in the Americas, despite not being captured by the international community. To investigate whether Zika virus transmission is still ongoing, we used international travel data to reveal that local outbreaks were still occurring in 2017, despite relatively few cases being reported (**Fig. 1**). Our data demonstrate that the vast majority of Zika cases during 2017 were the result of an unreported Zika outbreak in Cuba, which occurred while public data suggested the epidemic was nearing its end in the Americas (PAHO, 2017a) (**Fig. 1**).

**Figure 1.**
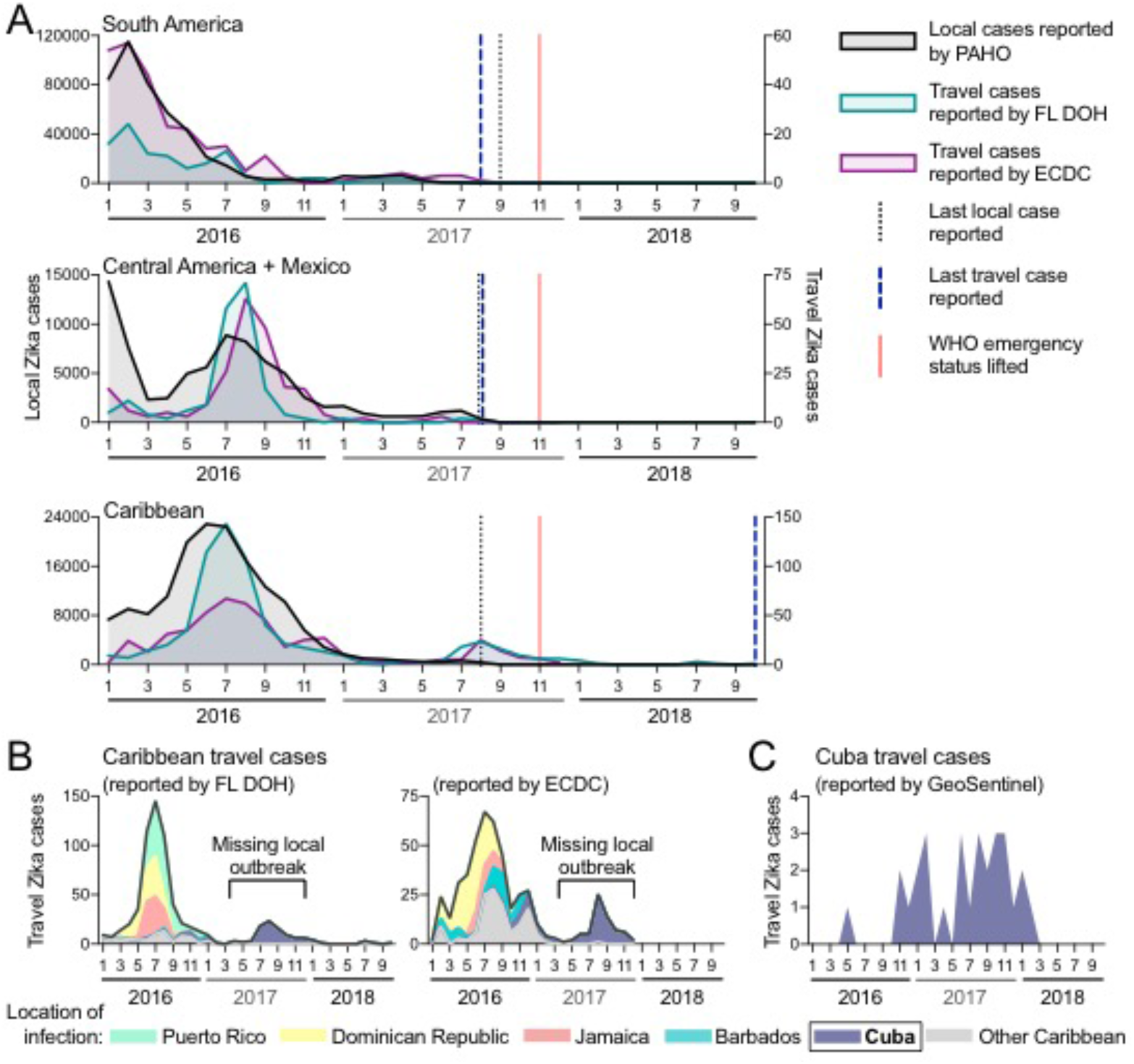
International travel cases reveal unreported Zika outbreak in Cuba in 2017. Local and travel-associated Zika cases were used to determine if outbreaks were still occurring during 2017. (**A**) Monthly local Zika cases (left y-axis) reported by the Pan American Health Organization (PAHO) and monthly travel-associated Zika cases (right y-axis) reported by the Florida Department of Health (FL-DOH) and the European Centre for Disease Prevention and Control (ECDC) were sorted by origin of exposure. The vertical lines represent the months the last local and travel cases were reported per region, and the month that the World Health Organization (WHO) Public Health Emergency of International Concern status was lifted for the Zika epidemic (November, 2017). In each region, travel cases and local cases were correlated (Pearson r range = 0.542-0.976, each comparison can be found in **Supplemental File 1**). (**B**) The total number of Zika cases reported by the FL-DOH and the ECDC associated with travel originating in the Caribbean are shown (black line) and are shaded by the top 5 origin locations (all other placed in the ‘Other Caribbean’ category). (**C**) Zika cases associated with travel from Cuba, diagnosed by the GeoSentinel Surveillance Network, were sorted by month of clinic visit. Travel cases diagnosed by the GeoSentinel Surveillance Network originating from other parts of the Americas are not shown. All of the data used for this figure can be found in **Supplemental File 1**.

To determine whether Zika case reports from international travelers could reveal outbreaks not captured by local case reports, we compared the temporal distribution of local and travel-associated Zika cases from 2016 to 2018 (**Fig. 1**). We obtained monthly suspected and confirmed Zika cases reported by individual countries and territories from PAHO. We obtained reports of international travel-associated Zika cases from the Florida Department of Health (FL-DOH) and the European Centre for Disease Prevention and Control (ECDC). We constructed Zika epidemic (epi) curves based on either local or travel-associated cases and found that they were in strong agreement from South America (Pearson *r* = 0.917 and 0.976 using FL-DOH and ECDC data, respectively) and the Caribbean (Pearson *r* = 0.828 and 0.856), and to a smaller extent Central America and Mexico (Pearson *r* = 0.542 and 0.583). For South America and Central America, we also found concordance for when the last local and travel cases were reported in August and September, 2017 (**Fig. 1A**).

We found that the last local case from the Caribbean was also reported in August, 2017. However, we observed a spike in Zika cases from travelers returning from the Caribbean during the summer of 2017 that were not captured by local reports, and Zika virus infected travelers from the Caribbean were reported until the end of the reporting period from the ECDC (December, 2017) and FL-DOH (October, 2018; **Fig. 1A**). By examining potential source locations for the travel-associated Zika cases in 2017, we found that between June, 2017 and October, 2018 more than 98% of them came from Cuba (90 of 91 Zika diagnoses in Florida, 63 of 64 Zika diagnoses in Europe; **Fig. 1B**). To further confirm the timing of a Zika outbreak in Cuba, we obtained travel-related infection data from the GeoSentinel Surveillance Network (Hamer et al., 2017; Leder et al., 2017) and found that 76% of the Zika cases associated with travel from Cuba were diagnosed in 2017 (22 of 29; **Fig. 1C**). While our travel data show that a Zika outbreak peaked in Cuba in 2017 with waning transmission continuing into 2018, during this time period no local Zika cases were reported by Cuba to PAHO or other international public health agencies (PAHO, 2017a).

### The Zika outbreak in Cuba was as large as those on other Caribbean islands

Using travel-associated Zika cases we identified an unreported Zika outbreak in Cuba that peaked during 2017 (**Fig. 1**). To investigate the size of the outbreak, we created a model using relationships between travel and local Zika case reporting, and found that it was likely as large as those on other Caribbean islands that peaked a year prior (**Fig. 2**).

In the absence of local case reporting, studies have demonstrated that travel-associated cases can be used to infer aspects of local virus transmission dynamics (Cauchemez et al., 2014; Fraser et al., 2009; Meltzer et al., 2008). Only 187 laboratory-confirmed Zika cases were reported by Cuba in 2016, and none were reported in 2017 (PAHO, 2017c). These reports are inconsistent with the outbreak dynamics that we detected using travel surveillance (**Figs. 1B, 2A**). To estimate the number of likely cases that should have been reported in Cuba in 2016 and 2017, we first investigated if travel cases accurately reflected the dynamics of known local Zika outbreaks for individual countries and territories outside Cuba (**Fig 2A**). We found that in places with at least 20 travel-associated Zika cases reported (**Fig. S1**), epi curves constructed from travel incidences were generally in agreement with epi curves generated from local reporting (mean Pearson *r* = 0.769, range = 0.121-0.984; **Supplemental File 1**).

We next constructed a Bayesian model to approximate the size of the Zika outbreak in Cuba using the mean posterior estimates of the proportion of local to travel incidence from 23 countries throughout the Americas (**Fig. S2**), each individually multiplied by the mean posterior estimates of the Cuba travel incidence rate (**Fig. S3**). Taking the population size of Cuba into account, we estimated that 5,707 Zika cases (interquartile range: 1,071 to 22,611) likely went unreported in this country (**Fig. 2B**), with the majority of these cases (>99%) having occurred in 2017. Our results therefore suggest that the 2017 Zika outbreak in Cuba was similar in size to the known 2016 outbreaks in countries with similar population sizes, such as Haiti (3,103 reported cases), Dominican Republic (5,305 reported cases), and Jamaica (7,165 reported cases; **Fig. 2C**).

**Figure 2.**
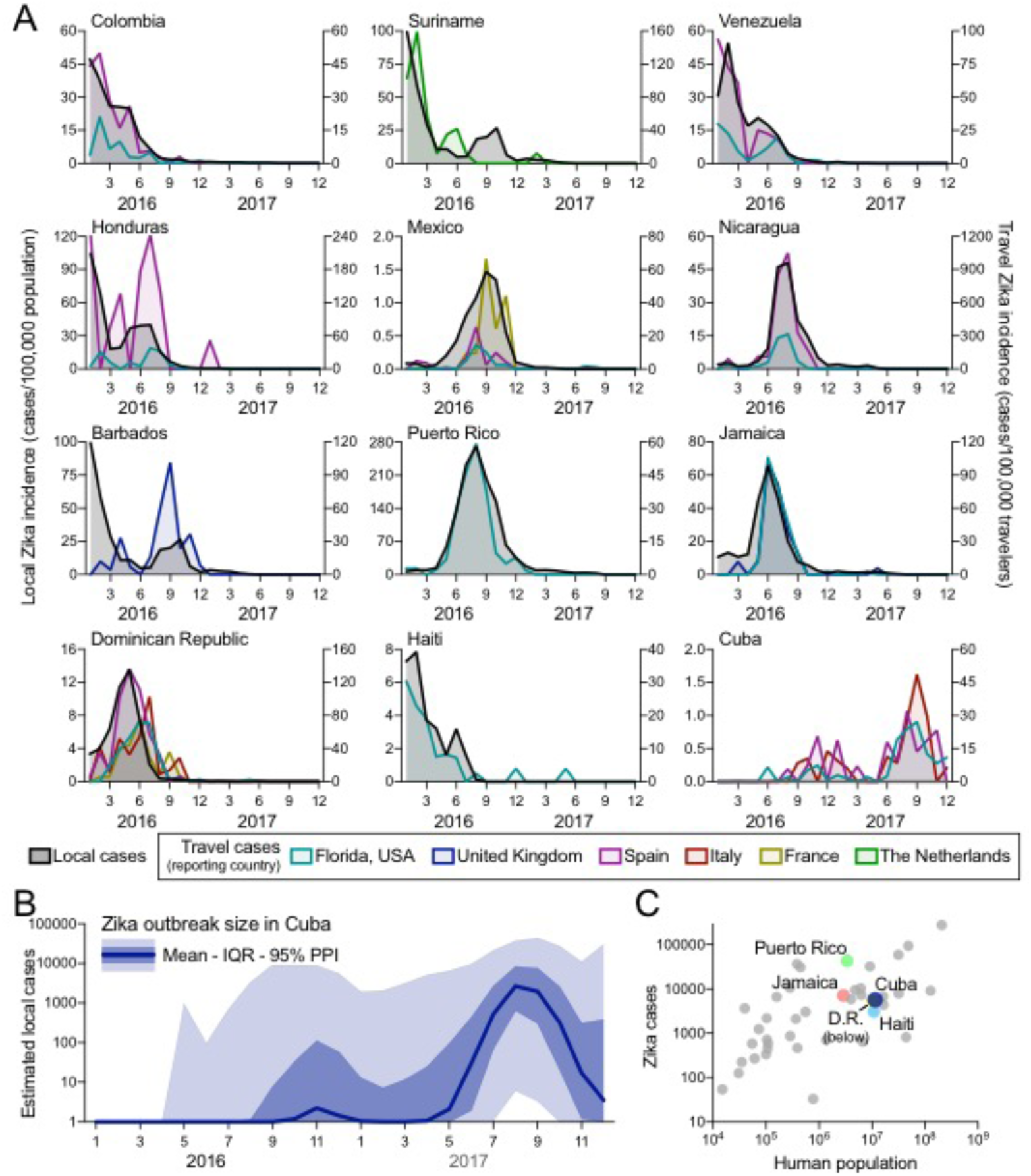
The Zika outbreak in Cuba during 2017 was similar in size to others during 2016. Infections of international travelers were used to estimate the size of the Zika outbreak in Cuba. (**A**) The local Zika virus incidence rates for each country/territory were calculated by the number of locally reported cases per month per 100,000 population. The travel Zika virus incidence rates for each country/territory of presumed exposure origin and reporting country (i.e. travel destination) pair were calculated by the number of travel-associated cases per month per 100,000 air passenger travelers entering the destination country from the origin. When there were at least 20 travel-associated Zika cases (**Fig. S1**), there was a positive correlation between travel and local incidence for all exposure origin and reporting country (i.e. travel destination) pairs (mean Pearson *r* = 0.769, range = 0.121-0.984; **Supplemental File 1**). (**B**) The number of Zika cases per month (mean, interquartile range, and 95% posterior predictive interval [PPI]) in Cuba during 2016–2017 were estimated by using fitted relationships between estimated local and travel incidence rates in countries with both sets of data to estimate what the local incidence rate in Cuba would have been if local data was available (**Figs. S2-S3**). This local incidence rate was then used to estimate local per capita incidence rates and subsequent number of Zika cases per month in Cuba. (**C**) The estimated number of Zika cases from Cuba (mean from **B**) and the total reported number of Zika cases during 2016–2017 from all countries/territories in the Americas with Zika virus transmission were plotted with the human population size from each region. Highlighted are the other large Caribbean countries/territories (D.R. = Dominican Republic.) All of the data used for this figure can be found in **Supplemental File 1**.

### Virus sequencing reveals multiple introductions of Zika virus into Cuba

We next determined the timing and origin of the Zika outbreak in Cuba using virus sequencing from travelers infected in Cuba. Our phylogenetic analysis showed that it was caused by multiple introductions of Zika virus from outbreaks in the Americas during the summer of 2016, corresponding to the season of peak *Ae. aegypti* transmission potential in Cuba (**Fig. 3**).

**Figure 3.**
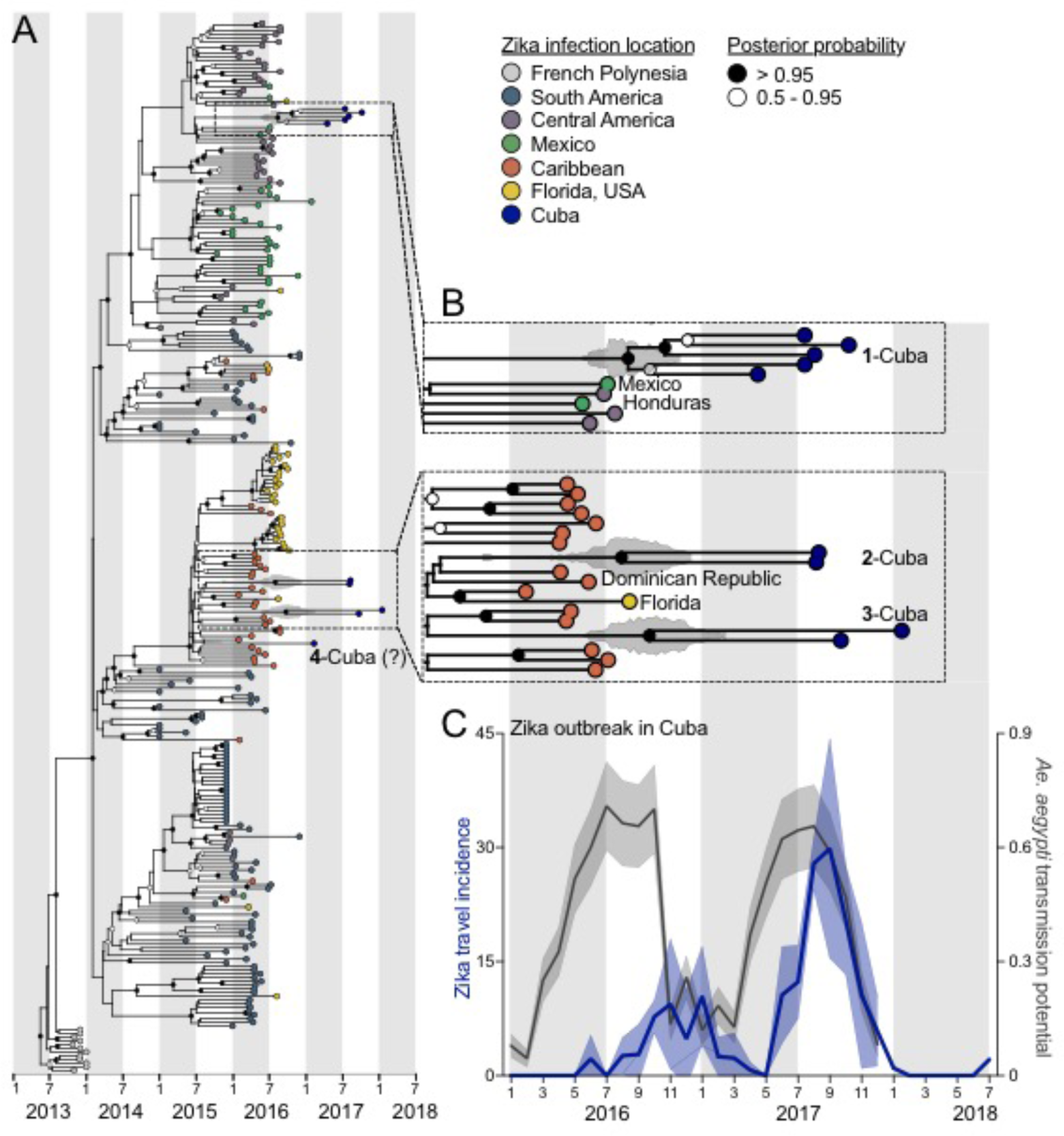
Multiple Zika virus introductions into Cuba during 2016 sparked the 2017 outbreak. Genomics approaches were used to determine the timing and sources of the Zika virus introductions into Cuba. (**A**) A maximum dade credibility tree was constructed using 283 near complete Zika virus protein coding sequences, including 10 sequences from travelers returning from Cuba during 2017–2018. (**B**) The zooms show the tMRCAs for each of the Cuba clades, as well as sequences basal on the tree. The fill color on each tip represents the probable location of infection, the clade posterior probabilities at each node are indicated by white circles tilled with black relative to the level of posterior support, and the grey violin plot indicates the 95% HPD interval for each tMRCA. The mean tMRCA for clade 1-Cuba was August, 2016 (95% HPD = May-November, 2016), the mean tMRCA for clade 2-Cuba was July, 2016 (95% HPD = March-December, 2016), and the mean tMRCA for clade 3-Cuba was September, 2016 (95% HPD = May, 2016-February, 2017). A maximum likelihood tree and a root-to-tip molecular clock are shown in **Fig. S4**. (**C**) The monthly Zika virus incidence rates (travel cases/100,000 travelers) associated with travel from Cuba (mean with standard deviation of travel cases reported from Florida, Spain, and Italy) and the monthly *Ae. aegypti* transmission potential (modeled using weather from Cuba and the optimal temperature ranges for mosquito-borne transmission, mean with standard deviation) were plotted to show their relationships to the Zika virus tMRCAs in Cuba. The data used to create panel b can be found in **Supplemental File 1**, and the Zika virus genomes sequenced during this study can be found in **Supplemental File 2**.

We sequenced Zika virus genomes from nine infected Florida travelers arriving from Cuba during 2017–2018 and obtained one Cuban Zika virus genome from GenBank (MF438286). In addition to our previous Zika virus sequences from the 2016 outbreak (Grubaugh et al., 2017), we also sequenced four additional genomes from Florida to demonstrate that the Zika virus lineages from Cuba were distinct from those in Florida, and thus *bonafide* travel-associated cases (**Fig. 3A**). We openly shared all sequences as they were generated (https://andersen-lab.com/secrets/data/zika-genomics/), and combined them with other publicly available sequences for a final dataset of 283 Zika virus genomes (**Supplemental File 2** and **Fig. 3**).

We constructed phylogenetic trees using time-resolved Bayesian inference (**Fig. 3A**) and maximum likelihood reconstruction (**Fig. S4**). We found that the Zika virus lineages in Cuba clustered with other virus genomes from the Americas, showing that the outbreak in Cuba was a continuation of the epidemic in the Americas, as opposed to introductions from ongoing Zika outbreaks in Asia (Lim et al., 2017; Watts et al., 2018) (**Fig. 3A**). Based on the placement of the Zika virus genomes from Cuba, we found evidence for one introduction from Central America (**Fig. 3B**, clade ‘1-Cuba’) and at least two from the Caribbean (**Fig. 3A, 3B**, clades ‘2-Cuba’, ‘3-Cuba’, and ‘4-Cuba’); however the placement of clade ‘4-Cuba’ is ambiguous, as it clusters with clade ‘3-Cuba’ in the maximum likelihood tree (**Fig. S4**), and may not be a seperate introduction (**Fig. S4**).

We estimated the time to the most recent common ancestor (tMRCA) for each of the Cuban Zika virus clades to be between July and September, 2016 (**Fig. 3B**). These findings suggest that the 2017 Zika outbreak in Cuba was caused by multiple introductions during the summer of 2016, corresponding to the peak of the Zika outbreaks in the Caribbean and Central America (June-September, 2016; **Fig. 1A**).

Zika virus is vectored by *Ae. aegypti*, so we next investigated if the introductions of the virus into Cuba aligned with the optimal time period for mosquito-borne transmission. Temperature is the primary seasonal factor driving Zika virus transmission as it influences mosquito development, survival, reproduction, biting rates, and vector competence (Caminade et al., 2016; Mordecai et al., 2017; Siraj et al., 2017). Thus, we used a model that estimated when transmission was most likely to occur based on favorable temperature ranges for mosquito-borne transmission (Mordecai et al., 2017). Using weather data for Cuba, we found that the Zika virus introductions into Cuba corresponded to the period of optimal local *Ae. aegypti* transmission potential (**Fig 3C**). The timing of multiple Zika virus introductions during the summer of 2016 is therefore unsurprising, as they appear to have occurred when Zika virus activity was peaking in source locations and local *Ae. aegypti* transmission potential in Cuba was high. These findings, however, do not explain why transmission in Cuba did not peak until 2017, which was at least a year later than other local outbreaks in the Americas (PAHO, 2017b) (**Fig. 2A**).

### Mosquito control may have delayed the Zika outbreak in Cuba

Given that Zika outbreaks were occurring throughout the Caribbean in 2016, and that the virus was introduced into Cuba in mid-2016, we investigated what factors may have been responsible for delaying the peak of the Cuban outbreak by a year (**Fig. 1**). We explored three primary hypotheses: (***1***) fewer opportunities for Zika virus introductions into Cuba in 2015 when introductions were occurring elsewhere in the Caribbean (Faria et al., 2017; Metsky et al., 2017), (***2***) differences in environmental conditions in 2016 lowering the susceptibility for *Ae. aegypti-borne* virus outbreaks, and (***3***) *Ae. aegypti* surveillance and control campaigns (Gorry, 2016; Reardon, 2016) limiting virus transmission. To investigate the likelihood of each hypothesis, we examined international travel patterns, yearly transmission of dengue virus (also vectored by *Ae. aegypti)*, local weather conditions, and news reports. Comparing all three hypotheses, we found that conditions in Cuba could likely have supported a large Zika outbreak in 2016, but that it may have been delayed by a country-wide *Ae. aegypti* control campaign (**Fig. 4**).

Outbreaks of *Ae. aegypti-borne* viruses, including Zika virus, require conducive conditions to support sustained transmission and opportunities for virus introductions. As air travel is the main source of long-distance virus dispersion (Khan et al., 2014; Nunes et al., 2014; Semenza et al., 2014), we analyzed air travel patterns to determine if Cuba had fewer opportunities for virus introductions early during the epidemic, potentially delaying the outbreak (**Fig. 4A**). Using monthly airline passenger arrivals coming from all 48 countries and territories in the Americas known to have local Zika virus transmission from 2014–2017, we did not detect any large deviations in air traffic to Cuba when

**Figure 4.**
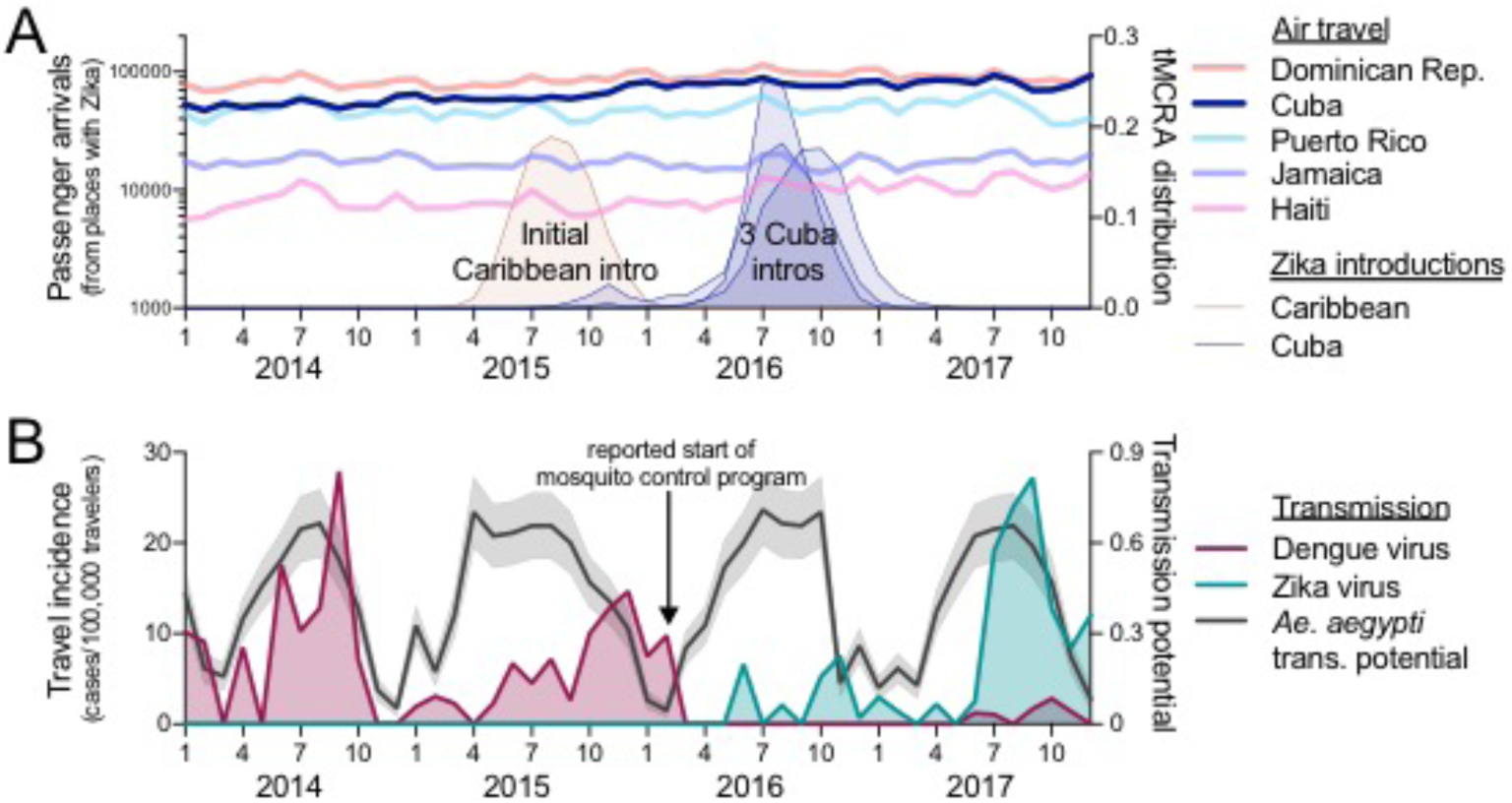
Aggressive *Aedes aegypti* control may have delayed Zika outbreak in Cuba. (**A**) The potential for Zika virus introductions was assessed by total airline passenger arrivals per month from 2014–2017 coming from regions in the Americas known to support local Zika virus transmission (excluding the continental United States because the outbreaks were relatively small), along the distribution of likely introduction tames (i.e. tMRCAs) of the initial (known) Zika virus introduction in the Caribbean (tMRCA April - December, 2015) and the three separate introductions into Cuba (tMRCAs March, 2016 - February, 2017). Clade ‘Cuba-4’ has a “?” because the genome dusters with clade ‘Cuba-3) in the maximum likelihood tree (**Fig. S4**) and it may not represent a separate introduction. (**B**) Analysis of dengue and Zika virus incidence, *Ae. aegypti* transmission potential, and the timing of a reported vector control campaign were used to investigate the delayed Zika outbreak in Cuba. Monthly dengue and Zika virus travel incidence rates (travel cases/100,000 travelers), as reported by the FL-DOH, and relative *Ae. aegypti*-borne virus transmission potential, determined by a temperature-sensitive model (Mordecai et al., 2017) and monthly temperature from Havana, Cuba, were used to fudge the impact of the aggressive *Ae. aegypti* mosquito control program that was reported to have begun in Cuba during February, 2016. The data used for this figure can be found in **Supplemental File 1**.

Zika virus was being introduced elsewhere in the Caribbean (starting in mid-2015; **Fig. 4A**) (Faria et al., 2017; Metsky et al., 2017). Moreover, air travel volumes were higher into Cuba than neighboring islands with large outbreaks in 2016, including Puerto Rico and Jamaica (**Fig. 4A**). These findings suggest that changes in air travel did not play a role in delaying the outbreak in Cuba.

It is possible that environmental conditions in Cuba in 2016 were not conducive for large *Ae. aegypti-*borne virus outbreaks. To explore this scenario, we examined transmission of another *Ae. aegypti-*borne virus, dengue virus, using incidence rates from travel-associated dengue cases reported by the FL-DOH (**Fig. 4B**). We found that following large dengue outbreaks in Cuba in 2014 and 2015, dengue virus transmission subsided in 2016 before increasing in 2017 (**Fig. 4B**). The dengue epi curves were similar to the Zika epi curves from 2016 to 2017 (**Fig. 4B**), supporting the hypothesis that conditions in Cuba were not conducive for large *Ae. aegypti-*borne virus outbreaks in 2016. These data, however, do not reveal if this was due to local weather conditions or human intervention.

As temperature is the primary driver of *Ae. aegypti-*borne virus transmission (Caminade et al., 2016; Mordecai et al., 2017; Siraj et al., 2017), we used our temperature-dependent transmission model (**Fig. 3C**) to determine if local weather in 2016 may have delayed the Zika outbreak in Cuba to 2017 (**Fig. 4B**). We found that this was likely not the case, as *Ae. aegypti* transmission potential was as high in 2016 as it was during the dengue outbreaks in 2014 and 2015, and the Zika outbreak in 2017 (**Fig. 4B**). These findings suggest that environmental factors were likely not responsible for delaying the Zika outbreak in Cuba.

We previously demonstrated that mosquito control campaigns can reduce *Ae. aegypti* populations and human Zika virus infections (Grubaugh et al., 2017). Cuba has a long history of successful *Ae. aegypti* control (Gubler, 1989; Toledo et al., 2007), and following the detection of the Zika outbreak in Brazil, the country implemented a “National Zika Action Plan” for aggressive *Ae. aegypti* mosquito surveillance and control (Gorry, 2016; Reardon, 2016). To investigate if mosquito control may have played a role in delaying the Zika outbreak in Cuba, we compared the reported start of the mosquito control campaign to Zika and dengue virus transmission (based on travel incidence rates) in Cuba (**Fig. 4B**). We found that following the implementation of mosquito control in February, 2016, travel incidence data showed minimal transmission of both dengue and Zika viruses throughout the year (**Fig. 4B**). By searching news articles for Zika and dengue in Cuba from 2015–2018, we found that Cuban officials reported that the mosquito control program was successful, based on a near elimination of dengue and very few Zika cases (see **Supplemental File 3** for a timeline of selected news articles). However, no information was reported on the overall length of the campaign, and importantly, if it was still ongoing when the Zika outbreak intensified in 2017. The timing of the mosquito control campaign, followed by a decrease in both dengue and Zika cases (**Fig. 4B**) - despite high transmission potential (**Fig. 4B**) - suggests that mosquito control efforts may have been responsible for delaying the Zika outbreak in Cuba. The outbreak the following year was likely preceded by a resurgence in *Ae. aegypti* populations, leading to an increase in transmission of Zika virus lineages that were introduced in 2016 (**Fig. 3**).

### Potential for global spread from unrecognized local outbreaks

Unrecognized and delayed outbreaks have the risk of ‘silently’ spreading viruses to other parts of the world. To investigate potential risks of undetected Zika outbreaks, we analyzed global air travel patterns and *Ae. aegypti* suitability, and identified several regions where Zika virus could have been introduced from an unrecognized outbreak in Cuba during 2017.

Based on the occurrence of travel-associated Zika cases reported by the FL-DOH and the ECDC, we found that Zika virus transmission in Cuba was the most intense between June-December of 2017 (**Fig. 5A**). We then used this time period to assess where local mosquito-borne Zika virus transmission could have been introduced from Cuba using global air travel data from Cuba and previously estimated world-wide *Ae. aegypti* suitability (Kraemer et al., 2015) (**Fig. 5B**). Out of a total of ~4 million air travelers departing Cuba between June and December of 2017, we found 18 countries and US states that received >20,000 travelers, with >100,000 arriving in Florida, Canada, Mexico, and Spain (**Fig. 5B**). Based on environmental suitability for *Ae. aegypti* of the 18 areas with >20,000 travelers from Cuba, we estimated that Florida, Mexico, Panama, Venezuela, and Colombia were most at risk of Zika virus having been introduced from Cuba during June-December, 2017 (Fig. 5B). Indeed, four local Zika cases were reported in Florida during 2017 linked to their partners having recently returned from Cuba (FL DOH, 2017a, 2017b, 2018). Despite these findings, however, beyond a few cases, no Zika outbreaks were reported in these 18 regions in 2017, perhaps due to existing herd immunity (Netto et al., 2017; Zambrana et al., 2018). These results show the global connectedness of Cuba, and with Zika cases associated with travel to Cuba still ongoing as of October, 2018 (**Fig. 1**), continued surveillance is required to detect potential further spread.

**Figure 5.**
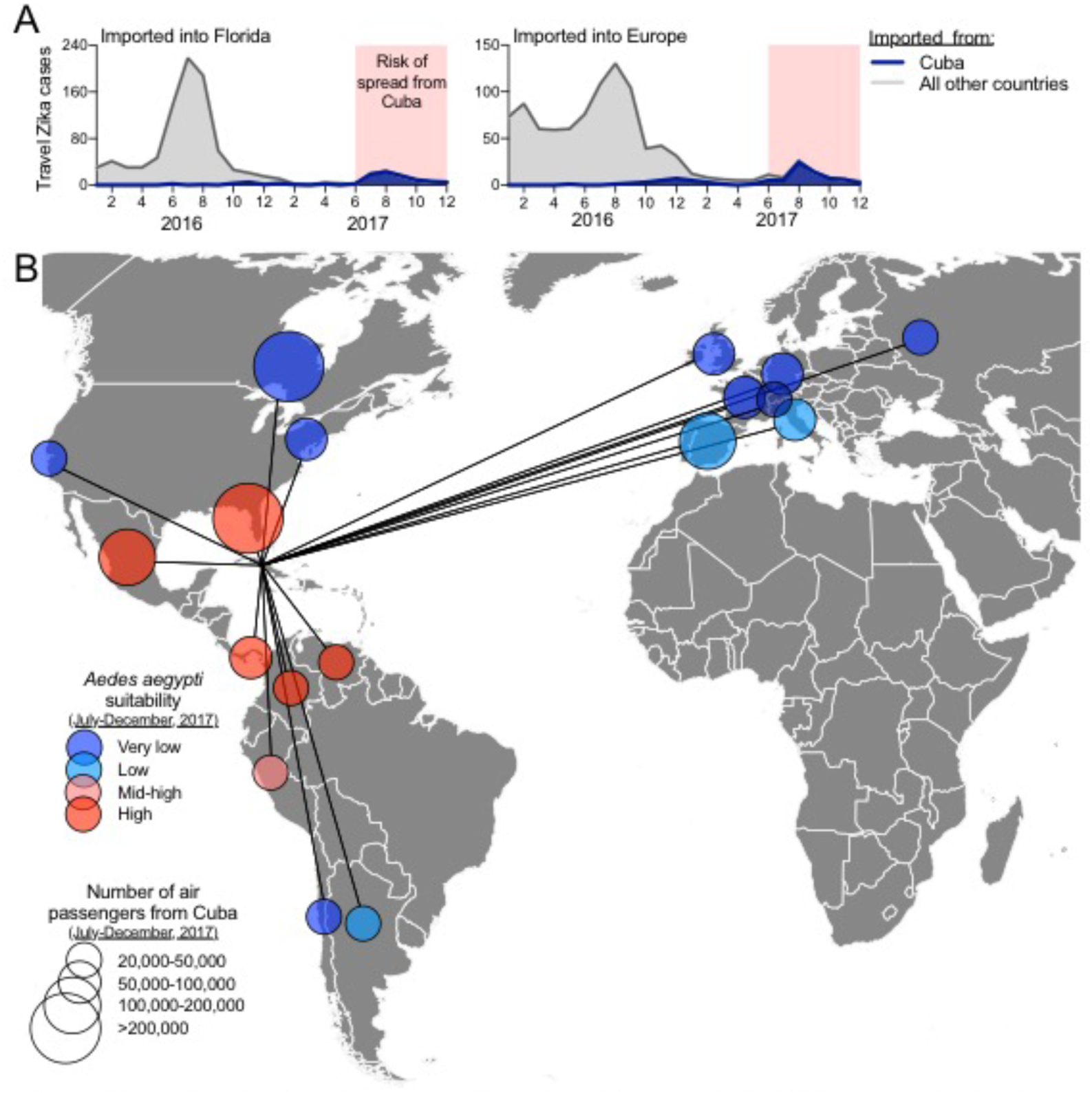
Risk of ‘silent’ Zika virus spread from the outbreak in Cuba during 2017. Travel volumes from Cuba and *Ae. aegypti* suitability were used to address the potential spread of Zika virus from Cuba during the outbreak in 2017. **(A)** Monthly Zika cases associated with international travel reported by the FL-DOH and the ECDC, sorted by travel origins in Cuba oral other countries/territories in the Americas, were used to demonstrate that >98% of all travel-associated Zika cases during June-December of 2017 came from Cuba. **(B)** During June-December, 2017, all countries and U.S. states that received >20,000 airline passengers from Cuba are shown, along with the relative *Ae*. *aegypti* suitability, to represent possible destinations for Zika virus spread from Cuba. The data used for this figure can be found in **Supplemental File 1**.

## Discussion

### Travel data to detect Zika outbreaks

Using travel data and virus genomics, we discovered a Zika outbreak in Cuba during 2017, a period in which the epidemic was waning across the Americas (PAHO, 2017a) (**Figs. 1 & 2**). Reports of an outbreak in Cuba did make the news in 2017 (Reuters, 2017), but critically, the outbreak was not reported by PAHO (PAHO, 2017a), or other public health agencies, and thus went largely undetected by the international community. With Zika outbreaks still arising in new locations, including Angola (Virological, 2018) and India (Pulla, 2018), both with possible origins in the Americas, it is important to identify and report lingering outbreaks to better prepare for potential future spread (Bogoch et al., 2016; Kraemer et al., 2017) (**Fig. 5**).

Epidemiological updates by the WHO and PAHO are the primary methods for disseminating information about infectious disease outbreaks and epidemics. Critically, they rely on accurate case reporting from individual countries and territories, but depending on resources and priorities, the reporting of local outbreaks may not be accurate. In this study, we investigated how Zika surveillance of international travelers can be used in conjunction with existing systems to fill knowledge gaps about ongoing outbreaks from places with irregular reporting that are difficult to sample. Our approach is particularly appropriate for regions such as the Caribbean, which, despite its long history of mosquito-borne virus outbreaks (Brathwaite Dick et al., 2012; Patterson et al., 2016; Weaver et al., 2018), is often understudied.

Using travelers for surveillance, however, is limited to where people primarily travel, which will be largely different for each destination. By using travel data from Europe, we were able to capture Zika cases from countries that we could not from Florida (**Fig. 2A**), but we did not detect any infected travelers coming from the Zika outbreaks in Angola (Virological, 2018) and India (Pulla, 2018) from any of our sources. Thus, using travelers as sentinels alone cannot provide a complete global picture of ongoing Zika outbreaks.

### Estimating the size of the Zika outbreak in Cuba

We estimate that the 2017 Zika outbreak in Cuba was similar in size to outbreaks from other Caribbean islands that peaked the year prior (**Fig. 2**). These analyses utilize the relationships between local- and travel-associated Zika data from non-Cuba countries, in combination with travel volumes and travel associated cases from Cuba. Other studies have used similar approaches to estimate the number of people infected with influenza A/H1N1 virus (Fraser et al., 2009) and the number of Middle East Respiratory Syndrome cases (Cauchemez et al., 2014). However, there are important limitations to such approaches that may influence our ability to estimate the size of the Zika outbreak in Cuba. First, accurate travel data are necessary to calculate travel incidence rates of Zika cases. This is challenging for Cuba, as travel policies from the United States have repeatedly changed during the past few years (Robles, 2016). To minimize this issue, we included air travel from both scheduled commercial flights from the International Air Transportation Association (IATA, 2018) and chartered flights from the U.S. Department of Transportation (US DOT, 2018). Additionally, we also obtained travel-associated Zika virus case data from Europe, where the travel policies to Cuba to the best of our knowledge have not recently changed.

Our estimated outbreak size also does not take into account differences in public health systems providing local data, and differing likelihoods of travelers becoming infected. For example, because of differences in public health infrastructure and resources, Zika case reporting in Haiti may be less accurate than Puerto Rico (Braga et al., 2017; Dowell et al., 2011). Additionally, exposures to mosquitoes are likely different between local residents and travelers, leading to differences in infection risks between the two populations. Such potential risk differences are largely unknown and dependent on location, behaviors, and length of stay (Cauchemez et al., 2014; Fraser et al., 2009), which could influence our estimates.

Finally, our size estimates are based on averaging across all regions, some of which may be more, or less, representative of the Zika outbreak in Cuba. While we found a strong correlation between epi curves generated from travel associated Zika cases and local reporting, variability among locations resulted in a wide interquartile range (1,071 to 22,611) on our mean estimate of 5,707 unreported Zika cases in Cuba. Our mean estimate, however, is consistent with the only two public reports from the outbreak in Cuba of 187 cases in 2016 reported by PAHO (PAHO, 2017c) and 1,847 cases in 2017 reported by the news agency Reuters (Reuters, 2017). Zika outbreaks from other locations in the Americas with comparable population sizes to Cuba were also reported to be similar in size (**Fig. 2C**).

### Multiple lineages of Zika virus ‘overwintering’ in Cuba

By sequencing Zika virus genomes from travelers infected in Cuba, we demonstrate that the 2017 outbreak was sparked by at least three introductions of the virus a year earlier (**Fig. 3**). Given our estimated size of the outbreak in Cuba, however, there are likely many additional Zika virus introductions not captured in our analyses (Grubaugh et al., 2017). Our tMRCA estimates of the Zika virus lineages from Cuba suggest that the virus survived the low mosquito abundance season (i.e. ‘overwintered’ from November to March) to cause more intense transmission in 2017 after the local mosquito population rebounded. While the factors supporting virus ‘overwintering’ are still unclear, it is plausible that Zika virus may have survived low mosquito abundance through a combination of low level mosquito-to-human transmission, vertical transmission in mosquitoes (da Costa et al., 2018; Thangamani et al., 2016), and, to a lesser extend, human sexual transmission (Althaus and Low, 2016). Considering that a large Zika outbreak in Cuba did not occur until after the viruses successfully ‘overwintered’, which may happen often with Zika outbreaks (Faria et al., 2017; Thézé et al., 2018), better understanding of how Zika virus is maintained when mosquito abundance is low might lead to novel control methods.

### Factors responsible for delaying the Zika outbreak in Cuba

By investigating news reports and modeling mosquito abundance, our study suggests that the Zika outbreak in Cuba may have been delayed by an *Ae. aegypti* control campaign (**Fig. 4**) (Gorry, 2016; Reardon, 2016). This accomplishment highlights the value of mosquito control for limiting transmission (Grubaugh et al., 2017), as Cuba may have been able to reduce the local burden of both dengue and Zika, despite otherwise conducive ecological conditions to support transmission of the viruses (**Fig. 4B**). However, we were unable to confirm if the mosquito control campaign was indeed successful, or what was specifically done, and for how long. Rather, we had to rely on temperature-dependent modeling of *Ae. aegypti*-borne transmission and local news reporting to test our hypothesis that the delayed outbreak was environmental- or mosquito control-dependent. Having access to empirical mosquito abundance data would have allowed us to assess year-to-year differences in transmission potential and to specifically test if *Ae. aegypti* populations were reduced during the control campaign (Grubaugh et al., 2017).

### Future projection

All available data suggest that the Zika epidemic is waning in the Americas (**Fig. 1**). This includes the Zika outbreak in Cuba where, based on our travel data, the outbreak was significantly smaller in 2018 as compared to 2017 (**Fig. 1B**). Cryptic Zika virus transmission is likely still occurring in regions of the Americas, however, incomplete surveillance and reporting make this difficult to confirm and quantify. Accurate Zika virus seroprevalence surveys (Balmaseda et al., 2017; Zambrana et al., 2018) are now needed to determine the extent of ‘hidden’ outbreaks of Zika, and to get more accurate measures of the true overall size of the epidemic. Open access to empirical mosquito abundance data is also critical for more precise forecasting of transmission potential and to evaluate control measures (Rund and Martinez, 2017); importantly, these efforts should be prioritized and more fully supported. Such initiatives, combined with our framework of using travelers as sentinels of Zika virus infections, can serve as complementary resources to detect, monitor, and reconstruct outbreaks when local surveillance is insufficient.

## Methods

### Ethical statement

This work was evaluated and approved by the Institutional Review Boards (IRB) at The Scripps Research Institute. This work was conducted as part of the public health response in Florida and samples were collected under a waiver of consent granted by the FL-DOH Human Research Protection Program. The work received a non-human subjects research designation (category 4 exemption) by the FL-DOH because this research was performed with remnant clinical diagnostic specimens involving no more than minimal risk. All samples were de-identified before receipt by the study investigators.

### Local Zika cases and incidence rates

PAHO is the primary source for information regarding Zika virus spread in the Americas, as well as suspected and confirmed cases per country and territory (PAHO, 2017a). Weekly case counts, however, are made available as cumulative cases, not the number of new cases per week. These data are often problematic for reconstructing outbreak dynamics because of reporting delays and ‘spikes’ (e.g. more than one week of cases submitted after weeks of no reporting). Curated weekly case counts per country and territory are presented as bar graphs (not as datasheets) (PAHO, 2017a). Therefore, to increase the accuracy of calculating Zika virus incidence rates, we captured screenshots of the 2016–2017 weekly Zika virus case (suspected and confirmed) distributions, and extracted the case counts using Web Plot Digitizer v3.10 (http://arohatgi.info/WebPlotDigitizer), which we previously validated (Grubaugh et al., 2017). Extracted case numbers were recorded in .csv files and aggregated per month for this analysis. Yearly human population numbers were retrieved from the United Nations Population Division (https://population.un.org/wpp/) and were used to calculate monthly local Zika virus incidence rates (suspected and confirmed Zika cases/100,000 population) per country and territory. Monthly Zika cases and incidence rates are available at: https://github.com/andersen-lab/paper_2018_cuba-travelzika.

### Travel-associated Zika and dengue cases and incidence rates

Weekly cumulative travel-associated Zika and dengue case numbers were collected from 2014–2018, and are publically available from the FL DOH (FL DOH, 2018). The cases reported on the FL DOH database include those that were confirmed by both PCR and serological assays, and within and without symptoms onset dates (note that many of the pregnant women that were serologically positive for Zika virus were asymptomatic). A travel history was also recorded for most patients. For this study, we only included PCR positive cases with a known date for the onset of symptoms and who only traveled to one international location within the 2 weeks prior to symptoms onset so we could more accurately sort the temporal and spatial distribution of travel-associated cases. We also excluded cases with sexual or congenital exposure. We aggregated the data by month of symptoms onset and by location of likely exposure (i.e. travel origin). Of the travel-associated Zika cases diagnosed in Florida (*n* = 1,333), 49% were visiting friends and relatives, 17% were refugees or immigrants, 17% were traveling for tourism, 3% were traveling for business, and 14% were traveling for unknown or other reasons. Of the travel-associated dengue virus cases where the questionnaire was given (only started for dengue in 2016, *n* = 88), 67% were visiting friends and relatives, 25% were traveling for tourism, and 8% were traveling for other reasons.

We also requested travel-associated Zika cases from the ECDC European Surveillance System (TESSy) (ECDC, 2017). We requested all travel-associated Zika cases reported to the ECDC during 2016–2017, sorted by month of symptoms onset, reporting country, and location of likely exposure. The data was provided by Austria, Belgium, Czech Republic, Denmark, Finland, France, Greece, Hungary, Ireland, Italy, Luxembourg, Malta, The Netherlands, Norway, Portugal, Romania, Slovakia, Slovenia, Spain, Sweden, and the United Kingdom, and released by ECDC. The raw travel-associated case counts from Europe has not been published, was obtained through specific request from the ECDC, and we do not have permission to make it public. In addition, the views and opinions that we expressed herein do not necessarily state or reflect those of ECDC. The accuracy of our statistical analysis and the findings we report are not the responsibility of ECDC. ECDC is not responsible for conclusions or opinions drawn from the data provided. ECDC is not responsible for the correctness of the data and for data management, data merging, and data collation after provision of the data. ECDC shall not be held liable for improper or incorrect use of the data.

Data on travelers to Cuba diagnosed at GeoSentinel sites were also analyzed. The GeoSentinel Global Surveillance Network consists of 72 specialized travel and tropical medicine clinics in 32 countries, and is staffed by specialists in travel and tropical medicine (http://www.istm.org/geosentinel). The GeoSentinel clinics provide routine clinical care to ill travelers and contribute de-identified demographic, travel, and clinical surveillance data on patients with travel-related illnesses to a centralized database (Harvey et al., 2013; Leder et al., 2013). Patient records with Cuba listed as the country of exposure and a diagnosis of mosquito-acquired Zika virus infection were extracted from the GeoSentinel database for the time period January 1, 2016 to November 12, 2018. Only confirmed cases were included in this analysis; these were defined as Zika virus PCR-positive in serum or urine, or Zika virus-specific IgM in serum and Zika virus antibody titers greater than four-fold higher than antibody titers for dengue or other flaviviruses or a four-fold rise in anti-Zika virus IgG and Zika virus antibody titers greater than four-fold higher than antibody titers for dengue or other flaviviruses (Hamer et al., 2017).

Monthly Zika virus travel incidence rates from all exposure (origin) and reporting (destination) combinations were calculated by number of travel-associated cases per 100,000 airline passengers (from origin to destination/month). Exposure-reporting combinations that accounted for less than 20 imported cases were not included in analysis. Air travel data was obtained as described below.

Though we previously hypothesized cruise ships may have an underrecognized role in Zika virus spread (Grubaugh et al., 2017), we did not use data from Zika virus infections that may have been associated with cruise travel, and thus did not collect cruise ship data for this study. First, there were very few infections linked to cruise travel in our dataset, which may be because these cases would more likely be tourists diagnosed elsewhere (and just visiting Florida for the cruise departure). Second, many of the reported cruise-related Zika infections were associated with more than one site for potential exposure, making it difficult to estimate local incidence rates (we removed all travel cases with multiple locations of potential exposure from our analyses). Third, scheduled cruise ship passengers arriving in Florida that stopped in Cuba are predicted to be substantially fewer (11,675/month scheduled for 2019; crawled from CruiseMapper: https://www.cruisemapper.com/) than air travel passengers from Cuba to Florida (80,366/month in 2017). Cruise travel between Cuba and Florida only began in 2016 (Vora, 2016), and thus there would have been even fewer passengers during our primary study period between 2016–2017.

The travel incidence rates derived from data collected from the FL DOH and ECDC and the curated travel-associated cases from Florida are available at: https://github.com/andersen-lab/paper_2018_cuba-travel-zika.

### Air passenger volumes

We collected air passenger volumes to calculate Zika and dengue virus travel incidence rates, to assess the potential for Zika virus importations into Cuba, and to investigate potential Zika virus spread from Cuba. From the IATA (IATA, 2018), we obtained the number of passengers traveling by air between all destinations in the Americas, plus to all global destinations from Cuba, from 2010–2017. IATA data consists of global ticket sales which account for true origins and final destinations, and represents 90% of all commercial flights. The remaining 10% of trips are modeled using airline market intelligence. One limitation of IATA data is it does not include chartered flights, which through our investigations, was only an issue for flights to and from the United States and Cuba. To make up for this, we obtained chartered flight data from Cuba to Florida during 2014–2017 from the US DOT (US DOT, 2018). The US DOT publically reports the number of passengers on all commercial and chartered flights departing and arriving in airports in the United States and includes origin and destination. Summarized air passenger volumes are available at: https://github.com/andersen-lab/paper_2018_cuba-travel-zika.

### Estimated local Zika cases in Cuba

We used two data types—locally acquired cases by country and Florida travel cases by country—to inform estimates of per capita local incidence in Cuba on a scale comparable to local incidence in other countries. We limited our analysis of countries besides Cuba to those with a correlation between monthly local and travel cases >0.25 (n=27), which appeared to be a natural breakpoint in the distribution of correlations. For each, we used the fda (https://cran.r-project.org/web/packages/fda/index.html) package in R to model per capita local incidence of Zika over time with univariate cubic B-spline functions with four knots per year for two years (2016–2017) described by parameters. We assumed that incidence among travelers from each country followed the same temporal pattern as local incidence but the two differed in magnitude by a factor, which could be due to differences in exposure or health-seeking behavior between international travelers and the general population. To estimate and for each of the 27 countries, we modeled local and travel incidence for each month as independent binomial random variables, with incidence as the number of “successes” and country population and number of travelers, respectively, as the number of “trials.” Logit-transformed values of the spline functions informed the probability of success in each trial. Based on this likelihood formulation and with non-informative priors, we estimated and for each country using a Metropolis-Hastings implementation of Markov chain Monte Carlo (MCMC). We assessed convergence by calculating Gelman-Rubin statistics on five replicate chains, and we performed posterior predictive checks on cumulative local incidence (**Fig. S2**) and travel incidence (**Fig. S3**) (Thompson Hobbs and Hooten, 2015). On the basis of Bayesian *p*-values < 0.05 on these posterior predictive checks, we removed four countries from subsequent analyses (leaving n=23 countries). To estimate per capita local incidence in Cuba, we first estimated for Cuba in a similar manner, but based on travel data only. We then took 10^4^ values of drawn randomly from the posteriors of pooled across 23 countries and multiplied them by random samples from the posterior of per capita travel incidence curves from Cuba to obtain a set of 10^4^ predictions of per capita local incidence curves for Cuba. R code and posterior samples are available at: https://github.com/andersen-lab/paper_2018_cuba-travel-zika.

### Zika virus sequencing

Zika virus RNA was sequenced using a highly multiplexed PCR approach, called PrimalSeq, that we previously described (Grubaugh et al., 2018b; Quick et al., 2017). Detailed protocols, including the primer scheme “ZIKV - Asia/America - 400bp” we used here to amplify Zika virus, can be found online (http://grubaughlab.com/open-science/amplicon-sequencing/ and https://andersen-lab.com/secrets/protocols/). In brief, virus RNA (2 μL) was reverse transcribed into cDNA using Invitrogen SuperScript IV VILO (20 μL reactions). Virus cDNA (2 μL) was amplified in 35 × ~400 bp fragments from two multiplexed PCR reactions using Q5 DNA High-fidelity Polymerase (New England Biolabs). Virus amplicons from the two multiplex PCR reactions were purified and combined (25 ng each) prior to library preparation. The libraries were prepared using the Kapa Hyper prep kit (Kapa Biosystems, following the vendor’s protocols but with ¼ of the recommended reagents) and NEXTflex Dual-Indexed DNA Barcodes (BIOO Scientific, diluted to 250 nM). Mag-Bind TotalPure NGS beads (Omega) were used for all purification steps. The libraries were quantified and quality-checked using the Qubit (Thermo Fisher) and Bioanalyzer (Agilent). Paired-end 250 nt reads were generated using the MiSeq V2 500 cycle kits (Illumina).

Our open source software package, iVar (Grubaugh et al., 2018b), was used to process the Zika virus sequencing data and call the consensus sequences. Source code and detailed documentation for iVar can be found at https://github.com/andersen-lab/ivar. In brief, BWA (Li and Durbin, 2009) was used to align the paired-end reads to a reference genome (GenBank KX087101). The primer sequences were trimmed from the reads using a BED file, with the primer positions, followed by quality trimming. The consensus sequence was called by the majority nucleotide at each position with >10× coverage. All alignments and consensus sequences were visually inspected using Geneious v9.1.5 (Kearse et al., 2012). The Zika virus sequences generated here can be found using the NCBI Bioproject PRJNA438510.

### Phylogenetic analyses

All available complete or near complete Zika virus genomes of the Asian genotype from the Pacific and the Americas were retrieved from GenBank in August, 2018. A total of 283 Zika virus genomes collected between 2013 and 2018 from Cuba (*n* = 10, including 9 generated in this study) and elsewhere from the Pacific and the Americas (*n* = 273, including 4 generated in this study from Florida, USA) were codon-aligned together using MAFFT (Katoh and Standley, 2013) and inspected manually.

To determine the temporal signal of the sequence dataset, a maximum likelihood (ML) phylogeny was first reconstructed with RAxML (Stamatakis, 2014) using the general time-reversible (GTR) nucleotide substitution model and gamma-distributed rates amongst sites (Guindon and Gascuel, 2003; Yang, 1994). Then, a correlation between root-to-tip genetic divergence and date of sampling was conducted in TempEst (Guindon and Gascuel, 2003; Rambaut et al., 2016; Yang, 1994). Time-scaled phylogenetic trees were reconstructed using the Bayesian phylogenetic inference framework available in BEAST v1.10.2 (Suchard et al., 2018). Accommodating phylogenetic uncertainty, we used an HKY+⌈4 nucleotide substitution model for each codon position, allowing for relative rates between these positions to be estimated, and an uncorrelated relaxed molecular clock model, with an underlying lognormal distribution (Drummond et al., 2006), a non-parametric demographic prior (Gill et al., 2013) and otherwise default priors in BEAUti v1.10.2 (Suchard et al., 2018). The MCMC analysis was run for 800 million iterations, sampling every 100,000th iteration, using the BEAGLE library v2.1.2 to accelerate computation (Ayres et al., 2012). MCMC performance was inspected for convergence and for sufficient sampling using Tracer v.1.7.1 (Rambaut et al., 2018). After discarding the first 200 million iterations as burn-in, virus diffusion over time and space was summarised using a maximum clade credibility (MCC) tree using TreeAnnotator (Suchard et al., 2018). Tree visualizations were generated with the Phylo (Talevich et al., 2012) module from Biopython and matplotlib (Hunter, 2007). Raw MAFFT codon alignment data, PhyML tree, BEAST XML file, and BEAST MCC time-structured phylogeny can be found at: https://github.com/andersen-lab/paper_2018_cuba-travel-zika.

### Monthly *Aedes aegypti* transmission potential in Cuba

Temperature is an important predictor of *Ae. aegypti-*borne virus transmission, as it affects mosquito population sizes (i.e. mosquito development, survival, and reproduction rates), interactions between mosquitoes and human hosts (i.e. biting rates), and mosquito transmission competence (i.e. mosquito infection and transmission rates) (Caminade et al., 2016; Mordecai et al., 2017; Siraj et al., 2017). Virus transmission by *Ae. aegypti* can occur between 18–34°C and peaks at 26–29°C (Mordecai et al., 2017). To assess yearly and seasonal variations in *Ae. aegypti* transmission potential for dengue and Zika virus, we used a temperature-dependent model of transmission using a previously developed R_0_ framework (Mordecai et al., 2017). By focusing this analysis on Havana, we controlled for spatial drivers of transmission and thereby isolated a representative example of temporal patterns in transmission potential. Using hourly temperature data obtained from OpenWeatherMap (https://openweathermap.org/), we calculated monthly mean temperature and used it to calculate monthly R_0_ as estimated by Mordecai et al. (Mordecai et al., 2017) (https://figshare.com/s/b79bc7537201e7b5603f). Doing so for 5,000 samples from the posterior of temperature-R_0_ relationships and normalizing between 0 and 1 yielded a description of relative *Ae. aegypti* transmission potential per month in Havana, Cuba during 2014–2017. Aggregated monthly weather data for and model outputs are available at: https://github.com/andersen-lab/paper_2018_cuba-travel-zika.

### Relative global *Aedes aegypti* suitability

To investigate the potential for Zika virus transmission and establishment, we used previously generated *Ae. aegypti* suitability maps (Kraemer et al., 2015) based on the statistical relationships between mosquito presence and environmental correlates (Bogoch et al., 2016). Maps were produced at a 5-km × 5-km resolution for each calendar month and then aggregated to the level of the U.S. states, countries, and territories, as used previously (Gardner et al., 2018). Relative *Ae. aegypti* suitability (i.e. very low, low, mid-high, and high) was then derived by using the mean aggregated values for each U.S. state, country, and territory, and also the mean value for the study period (June-December, 2017). The U.S. state, country, and territory suitability means and standard deviations can be found at: https://github.com/andersen-lab/paper_2018_cuba-travel-zika.

## Supporting information

## Data availability

The Zika virus sequences generated here can be found using the NCBI BioProject PRJNA438510. All data used to create the figures can be found in the supplemental files. The raw data and results for our analyses can be found at: https://github.com/andersen-lab/paper_2018_cuba-travel-zika.

## Acknowledgements

We thank R. Robles-Sikisaka, M.R. Wiley, K. Prieto, S. Taylor, and P. Jack for laboratory support and the ECDC for graciously providing data regarding travel-associated Zika virus infections. The findings and conclusions in this report are those of the authors and do not necessarily represent the official position of the US Army, CDC, ECDC, or the FL-DOH. RJO received support from the Arthur J. Schmitt Leadership Fellowship in Science and Engineering and from an Eck Institute for Global Health Graduate Fellowship. MUGK is supported by The Branco Weiss Fellowship - Society in Science, administered by the ETH Zurich and acknowledges funding from a Training Grant from the National Institute of Child Health and Human Development (T32HD040128). CBFV is supported by NWO Rubicon 019.181EN.004. GeoSentinel, the Global Surveillance Network of the International Society of Travel Medicine, is supported by a cooperative agreement (U50CK00189) from the Centers for Disease Control and Prevention, as well as the International Society of Travel Medicine, and the Public Health Agency of Canada. GB acknowledges support from the Interne Fondsen KU Leuven / Internal Funds KU Leuven under grant agreement C14/18/094. TAP received support from a RAPID award from the National Science Foundation (DEB 1641130) and a Young Faculty Award from the Defense Advanced Research Projects Agency (D16AP0014). SI and SFM are supported by NIH NIAID R01AI099210. KGA is a Pew Biomedical Scholar, and is supported by NIH NCATS CTSA UL1TR002550, NIAID contract HHSN272201400048C, NIAID R21AI137690, NIAID U19AI135995, and The Ray Thomas Foundation.

## GeoSentinel Contributors

Members of the GeoSentinel Surveillance Network who contributed data: Emmanuel Bottieau and Johannes Clerinx, Institute of Tropical Medicine (Antwerp, Belgium); Israel Molina, Hospital General Vall d’Hebron (Barcelona, Spain); Denis Malvy & Alexandre Duvignaud, Santé-Voyages et Médecine Tropicale (Bordeaux, France); François Chappuis and Gilles Eperon, Division of Tropical and Humanitarian Medicine – Geneva University Hospitals (Geneva, Switzerland); Sabine Jordan, University Medical Centre Hamburg Eppendorf, Division of Infectious Diseases and Tropical Medicine (Hamburg, Germany); Eli Schwartz, Sheba Medical Center (Tel HaShomer, Israel); Marta Díaz-Menéndez & Clara Crespillo-Andújar, Hospital La Paz-Carlos III (Madrid, Spain); Rogelio Lopez-Velez & Francesca Norman, Hospital Ramon y Cajal (Madrid, Spain); Kunjana Mavunda, International Travel Clinic (Miami, Florida, USA); Frank von Sonnenburg, Peter Sothmann & Camilla Rothe, University of Munich - Dept. of Infectious Diseases and Tropical Medicine (Munich, Germany); Federico Gobbi, Centre for Tropical Diseases, Ospedale Sacro-Cuore Don Calabria (Negrar, Italy); Anne McCarthy, Division of Infectious Diseases - Ottawa Hospital General Campus (Ottawa, Canada); Eric Caumes, Service des Maladies Infectieuses et Tropicales - Hôpital Pitié-Salpêtrière, Sorbonne Université (Paris, France); Hilmir Asgeirsson & Hedvig Glans (Clinic of Infectious Diseases - Karolinska University Hospital); Yukihiro Yoshimura, Department of Infectious Diseases - Yokohama Municipal Citizen’s Hospital (Yokohama, Japan).

## Author contributions

All contributions are listed in order of authorship. Designed the experiments: N.D.G, L.M.G., T.A.P., G.B., K.K., A.M., S.S., S.I., S.F.M., and K.G.A. Provided samples or data: N.D.G., S.S., K.G., A.W., A.L.T., J.T.L., M.U.G.K., A.H., D.B., D.S., B.S., V.L., I.S., M.R.C., E.W.K., A.C.C., L.H-L., S.W., L.D.G., M.J.R., J.K., P.K.L., D.M.M., D.I.W., G.P., D.H.H., GeoSentinel Contributors, K.K., A.M., S.I., and S.F.M. Performed the sequencing: N.D.G., S.S., G.O., N.L.M., M.R.W., and K.P. Analyzed the data: N.D.G., S.S., K.G., R.J.O., J.T.L., C.B.F.V., T.A.P., G.B., and K.G.A. Wrote manuscript: N.D.G., S.S., and K.G.A. Edited manuscript: K.G., R.J.O., M.U.G.K, C.B.F.V., L.D.G., L.M.G., T.A.P., G.B., A.M., S.I., and S.F.M. All authors read and approved the manuscript.

## Supplemental files

**Supplemental File 1**. Data used to create each figure.

**Supplemental File 2**. Zika virus genomes generated during this study.

**Supplemental File 3**. Timeline of selected news articles from 2015–2017 related to Zika and dengue virus transmission in Cuba.

**Supplemental File 4**. Raw MAFFT codon alignment data, PhyML tree, and BEAST MCC time-structured phylogeny.

---

**Figure S1.**
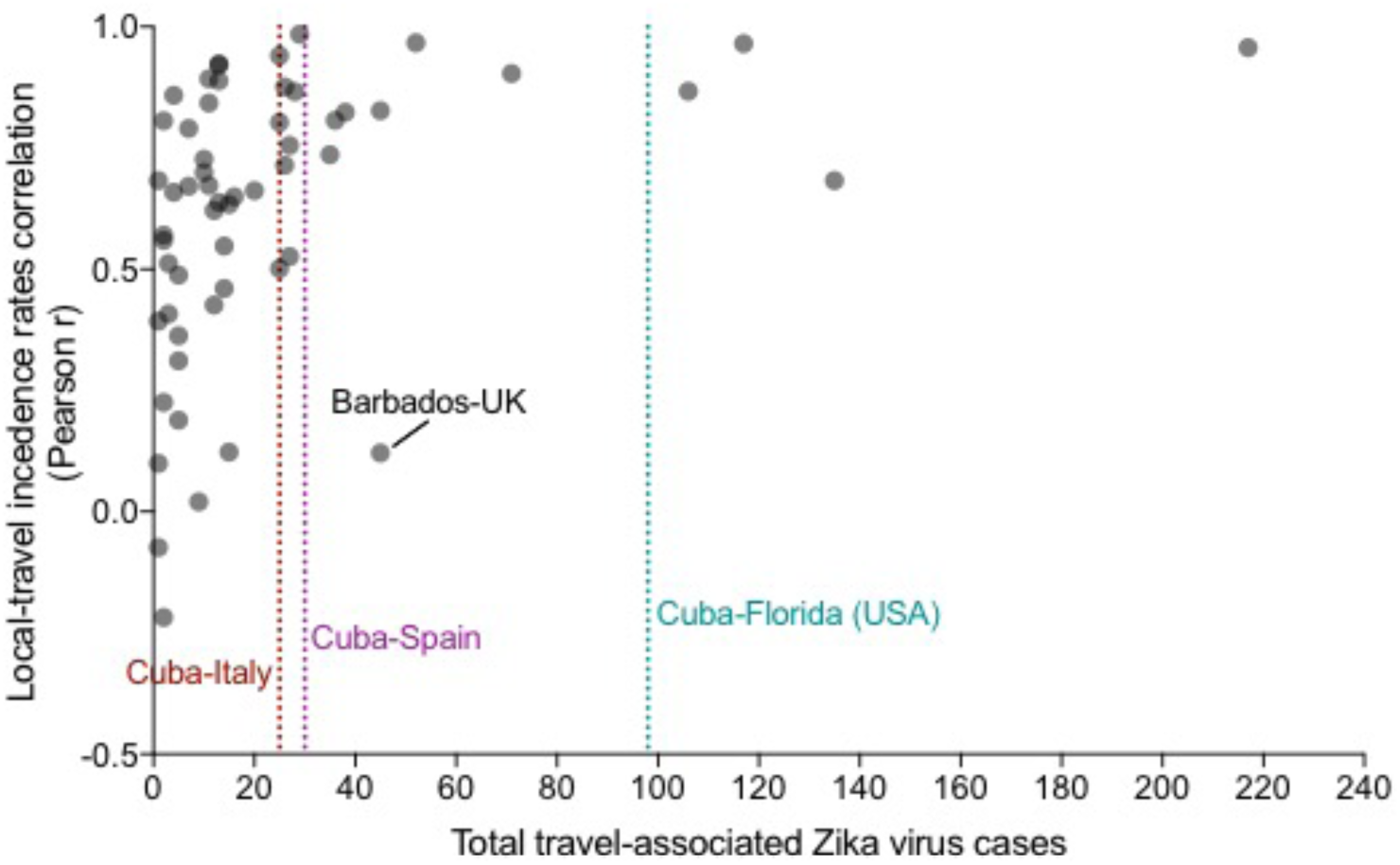
Relationship between the number of travel-associated Zika cases and the correlation between local and travel incidence rates. To determine the number of travel-associated infections needed to infer the shape of a local outbreak, we compared the total travel-associated Zika cases from each exposure-reporting country/territory combination (x-axis) with Pearson correlation between the local and travel incidence rates corresponding to the combination (y-axis). The travel-associated Zika cases were totaled from 2016–2017. For the Pearson correlations between local-travel incidence rates, monthly incidence values from 2016–2017 were compared. When there were >20 travel-associated cases, the local-travel Pearson *r* was >0.5, indicating a strong positive correlation and that the travel cases can help determine the shape of the local outbreak. The lone exception to that finding was from travelers from Barbados diagnosed in the United Kingdom (UK) because the travel cases miss the locally reported Zika virus peak during January-February, 2016, but they correlate with the second local peak during July-October, 2016 (**Fig. 2A**). In our dataset, there were 25 Zika virus infections diagnosed in Italy with recent travel to Cuba, 30 diagnosed in Spain, and 98 diagnosed in Florida. These totals are all within the range of strong positive correlations between local and travel incidence, justifying their use to infer the local Zika outbreak dynamics in Cuba (**Fig. 2B**).

**Figure S2.**
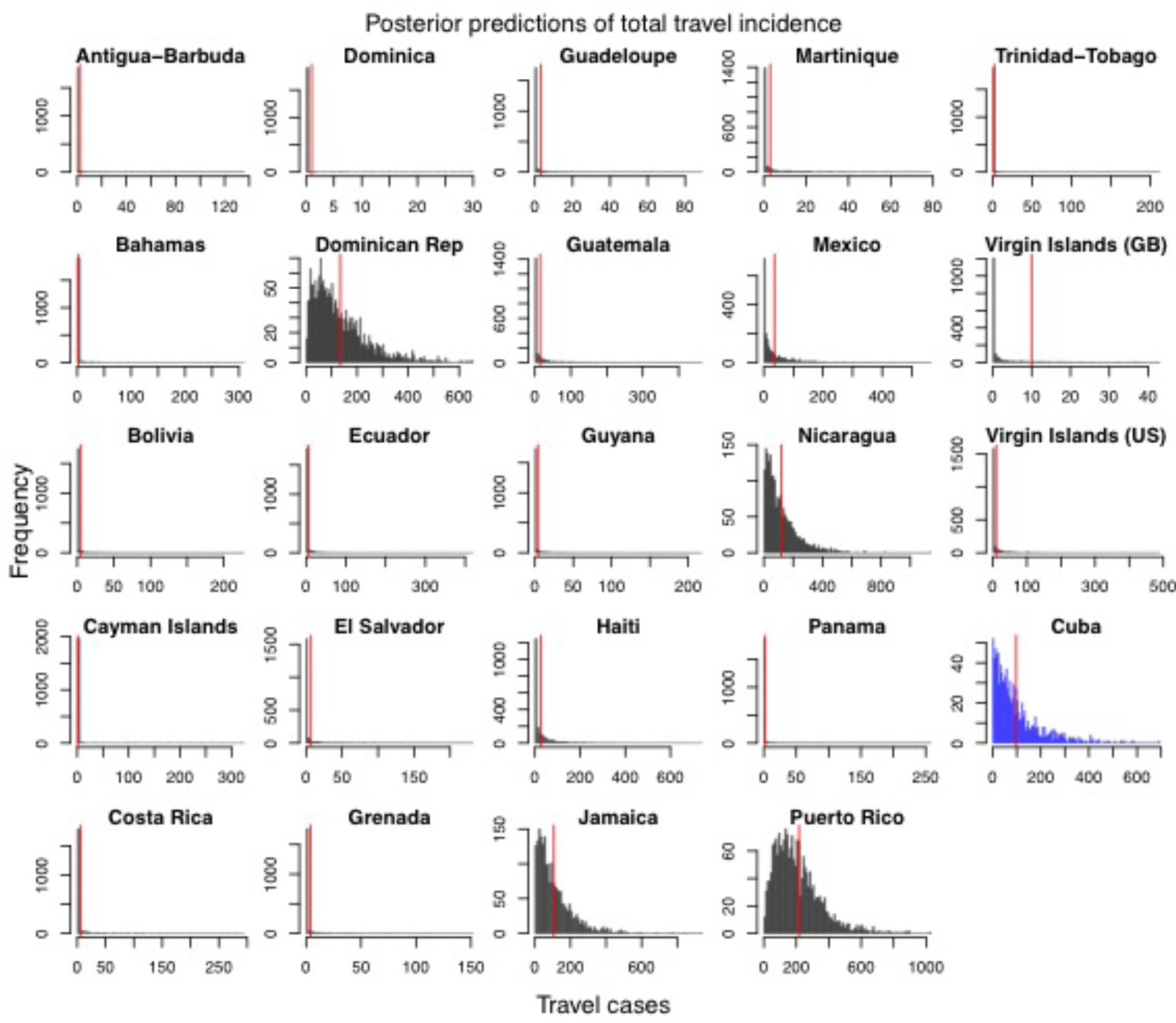
Posterior predictions of estimated total travel incidence from origin country into Florida. These distributions were used to inform the joint distribution between travel incidence and local incidence that was used to estimate local incidence in Cuba. Empirical total travel cases per country indicated by red vertical line. All of the countries shown above, besides Cuba, had >0.25 correlation between local incidence and travel incidence and had the observed value fall within the 95% posterior predictive interval of the distribution.

**Figure S3.**
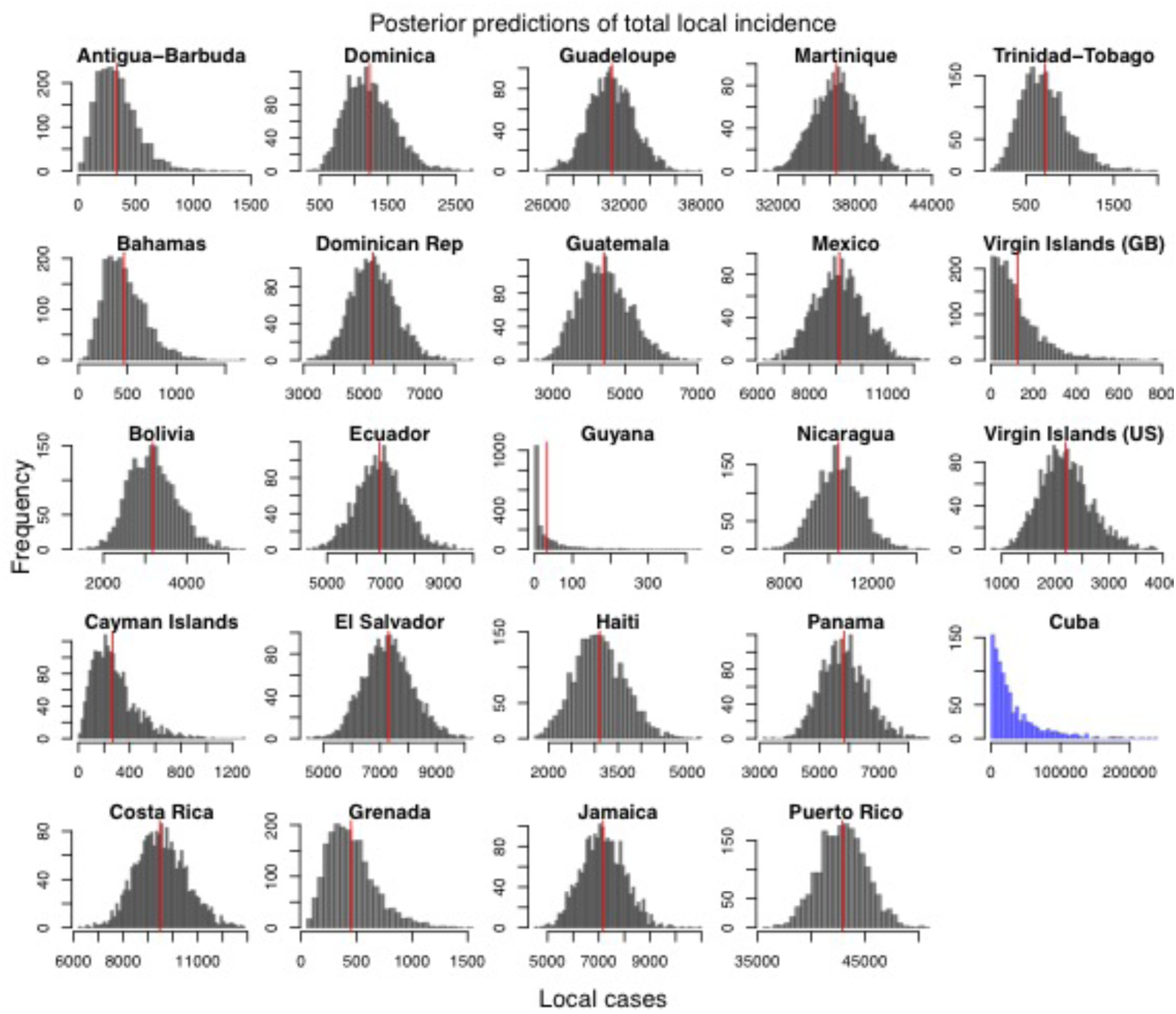
Posterior predictions of estimated total local incidence from origin country into Florida. These distributions were used to inform the joint distribution between travel incidence and local incidence that was used to estimate local incidence in Cuba. Empirical total local cases per country indicated by red vertical line. Estimated local incidence of Cuba indicated in blue with no empirical value. All of the countries shown above, besides Cuba, had >0.25 correlation between local incidence and travel incidence and had the observed value fall within the 95% posterior predictive interval of the distribution.

**Figure S4.**
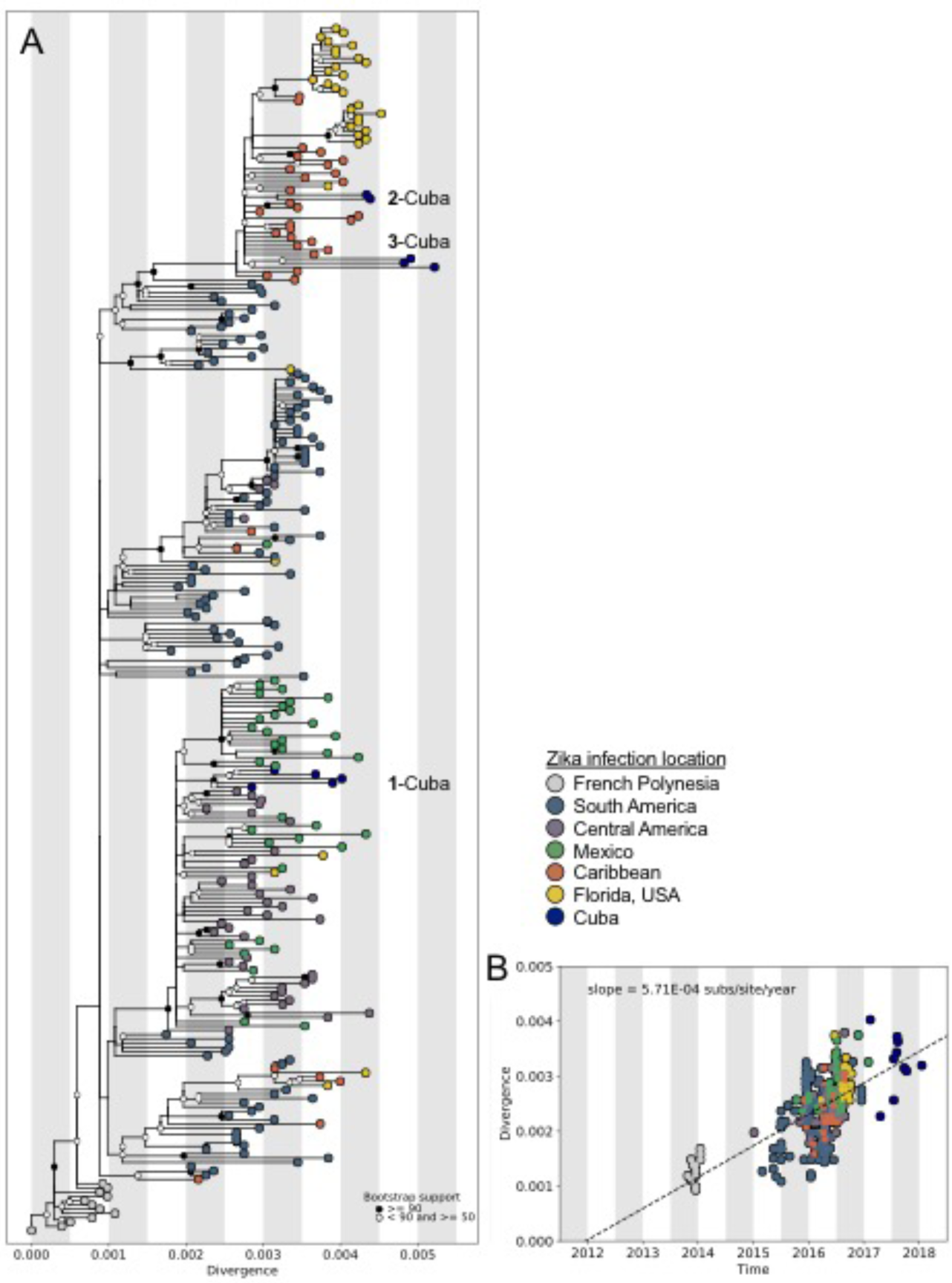
Maximum likelihood tree and root-to-tip regression of Zika virus genomes from Cuba and the epidemic in the Americas. (**A**) Maximum likelihood tree of publicly available Zika virus sequences (*n* = 269) and sequences generated in this study (*n* = 14). Tips are coloured by location. Bootstrap support values are colored at the nodes. Divergence shown as substitutions per site. “1-3 Cuba” represent three independent introductions of Zika virus into Cuba. (**B**) Linear regression of sample tip dates against divergence from root based on sequences with known collection dates estimates an evolutionary rate for the Zika virus phylogeny of 5.71 × 10^−3^ nucleotide substitutions per site per year.

