## Supplementary material for "International travelers and genomics uncover a ‘hidden’ Zika outbreak"

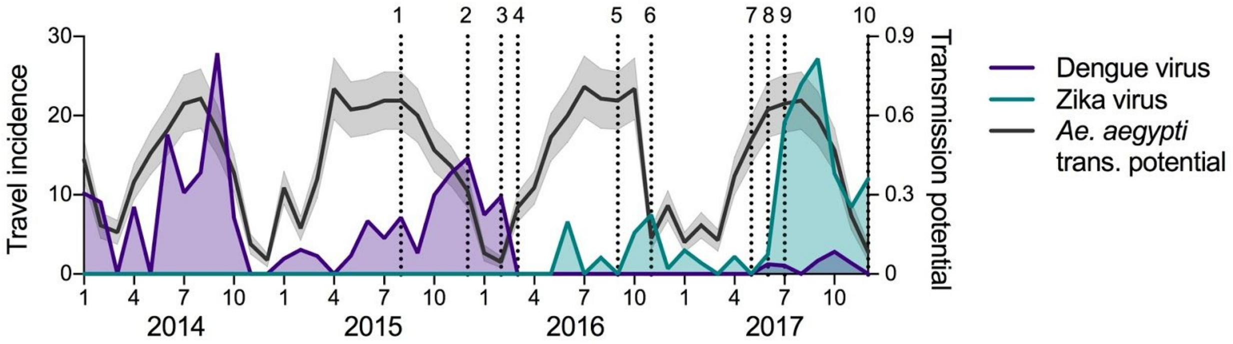


| # | Date | Highlights | Reference |
|---|---|---|---|
| 1 | 2015.08.07 | • Cuba reports high *Aedes aegypti* populations | https://havanatimes.org/?p=113113 |
| 2 | 2015.12.01 | • Cuba issues Zika epidemiological alert detailing surveillance, prevention, and control measures | https://scielosp.org/scielo.php?pid=S1555-79602016000100006&script=sci_arttext&tlng=pt |
| 3 | 2016.02.22 | • Cuba mobilizes 9,000 armed forces reserves and active members for mosquito fumigation | https://www.reuters.com/article/us-health-zika-cuba-idUSKCN0VV1EW |
| 4 | 2016.03.16 | • First autochthonous case diagnosed in Cuba | https://scielosp.org/scielo.php?pid=S1555-79602016000100006&script=sci_arttext&tlng=pt |
| 5 | 2016.09.28 | • No Zika cases reported since March<br>• Dengue nearly eliminated | https://www.reuters.com/article/us-health-zika-cuba-idUSKCN0ZE1TG |
| 6 | 2016.11.08 | • Zika outbreaks causing local transmission have occurred in Havana and two other cities | https://www.statnews.com/2016/11/08/zika-in-cuba/ |
| 7 | 2017.04.18 | • Zika cases spike around 1900 local cases | https://in.reuters.com/article/us-health-zika-cuba/cuba-says-zika-tally-rises-to-nearly-1900-cases-idINKCN18E25G |
| 8 | 2017.06.03 | • Reported increase *Aedes aegypti* populations in Cuba | https://www.telesurtv.net/english/news/Cuba-Fights-Rising-Dengue-Zika-Infestations-20170603-0015.html |
| 9 | 2017.07.11 | • Mosquito control program extended until late July | https://www.cubanet.org/mas-noticias/se-agrava-la-situacion-epidemiologica-en-holguin/ |
| 10 | 2017.12.26 | • Zika transmission reported in 18 municipalities<br>• Continued decline in Zika cases | http://en.escambray.cu/2017/cuba-reports-lowest-infant-mortality-rate-ever/ |
